# EngA adopts multiple distinct conformational states during ribosome assembly

**DOI:** 10.64898/2026.11.04.764183

**Authors:** Ankita Arora, Jhalak Aggarwal, Nisheeth Agarwal, Prem S. Kaushal

**Affiliations:** Laboratory of Ribosome Biology, UNECSO-DBT, Regional Centre for Biotechnology, NCR Biotech Science Cluster, Faridabad -121001, INDIA; Mycobacterial Genetics Laboratory, BRIC-Translational Health Science and Technology Institute, NCR Biotech Science Cluster, Faridabad-121001, INDIA

## Abstract

Ribosome assembly, a highly coordinated and crucial cellular process, is assisted by various assembly factors, including GTPases. EngA, an essential P-loop GTPase, is involved in late-stage assembly of the 50S ribosomal subunit. The structures of 50S ribosome in complex with EngA-bound GMPPNP, and EngA-bound GDP alone, without 50S subunit, have been reported previously. However, the molecular mechanism of EngA GTP hydrolysis-assisted 50S maturation remains poorly understood, and structures of the 50S ribosome in complex with EngA-bound GDP have yet to be deciphered. We have reported cryo-EM structures of the *Mycobacterium smegmatis* 50S ribosome in complex with EngA-bound GTPγS or GDP at 2.8 - 3.5 Å resolution. Structural analysis revealed that EngA and 50S subunit undergo a coordinated structural rearrangement, and an assembly intermediate is formed in which the stem-loop of H80 makes base-pair interactions with the junction of H89 and H90, forming a unique stem-loop-junction (SLJ) motif. EngA-bound GDP adopts distinctly different conformations in its ribosome-bound and unbound states. In summary, this study proposed a multi-state conformational model for EngA-assisted ribosome assembly.

## Introduction

Ribosomes, the ribonucleoprotein complexes of megadalton (mDa) size, decode information present in mRNA into proteins. The functional 70S ribosomes in prokaryotes are composed of a 30S small subunit (SSU) and a 50S large subunit (LSU). Assembly of ribosomal subunits is a multi-state dynamic process crucial for ensuring error-free translation. Any defect in ribosome biogenesis leads to severe physiological consequences such as ribosomopathies in humans (Narla & Ebert, 2010; Mills & Green, 2017) or growth arrest in microbes (Hassan *et al*, 2026). Pioneering work led by Nomura (Traub & Nomura, 1968) and Nierhaus (Nierhaus & Dohme, 1974) provided the first glimpse of ribosome subunit assembly by showing that an active ribosomal subunit could be reconstituted in vitro from their isolated components, ribosomal RNAs (rRNA) and ribosomal proteins (r-proteins). However, the requirement of non-physiological conditions, such as high salt and high temperature, for efficient in vitro reconstitution clearly suggested the need for additional factors to accomplish this process under physiological conditions (Mizushima & Nomura, 1970; Rohl & Nierhaus, 1982).

Ribosome assembly is a highly complex process assisted by various factors, including ATP-dependent helicases, chaperones, rRNA modification factors, and GTPases (Davis & Williamson, 2017; Shajani *et al*, 2011; Nomura, 1973; Kaczanowska & Rydén-Aulin, 2007; Wilson & Nierhaus, 2007). The ribosome assembly GTPases belong to the P-loop NTPase family, which is present across all three domains of life. These GTPases act as molecular switches, which alternate between an active ‘ON’ state and an inactive ‘OFF’ state. Such conformational changes are catalyzed by GTP binding and hydrolysis. In addition to ribosome biogenesis, these factors are involved in diverse cellular processes, such as signal transduction, translation, cytoskeletal formation, and intracellular transport. In bacteria, seven canonical GTPases (Era, RsgA, YchF, YihA, RbgA, ObgE and EngA) have been implicated in ribosome assembly to date (Britton, 2009; Karbstein, 2007; Caldon Pauline Yoong & March, 2001; Goto *et al*, 2013).

EngA, also known as Der in *Thermotoga maritima* (TmDer) and YphC in *Bacillus subtilis* (BsYphC), is an essential and highly conserved protein in eubacteria. (Verstraeten *et al*, 2011). It is structurally unique among all GTPases, possessing two consecutive N-terminal GTPase domains (GDs), GD1 and GD2, and a K-homology (KH) domain at the C-terminus (Hwang & Inouye, 2001). This distinctive architecture enables EngA to function as a multi-state molecular switch (Caldon Pauline Yoong & March, 2001), which predominantly associates with the 50S subunit during late assembly (Hwang & Inouye, 2006). Its depletion or mutation results in severe growth defects, altered polysome profiles, and the accumulation of immature 50S ribosomal subunits lacking ribosomal proteins, bL9, uL18, uL16, bL27, and bL36 (Hwang & Inouye, 2006)(Schaefer *et al*, 2006). EngA has also been shown to rescue defects in ribosome maturation resulting from loss of rRNA methylation at U2552, thereby indicating its role as an rRNA chaperone that stabilizes functional centers of the ribosome (Tan *et al*, 2002). The structural transition of EngA between active and inactive functional states strictly depends upon the occupancy of its nucleotide-binding pockets by GTP and GDP, respectively (Muench *et al*, 2006). EngA can also sense the intracellular GTP/GDP ratios, which affect its recruitment to the ribosome (Tomar *et al*, 2009). Furthermore, EngA functions synergistically with another assembly GTPase, RbgA, whereby prior association of RbgA with ribosomal intermediates increases the binding affinity of EngA (Arpin *et al*, 2025).

The crystal structures of GDP-bound EngA homologues TmDer and BsYphC show a compact cloverleaf-like arrangement, with the KH domain positioned between the two GTPase domains (Robinson *et al*, 2002; Muench *et al*, 2006). Conversely, Cryo-EM structures of 50S ribosome in complex with EngA-bound GMPPNP from *Escherichia coli* (EcEngA) at relatively low resolution ∼5.5 Å, and from *B. subtilius* BsYphC-GMPPNP at ∼3.3Å resolution, reveal a major domain rearrangement in EngA upon 50S binding, wherein the protein acquires an open conformation and binds to the inter-subunit face of the 50S subunit(Zhang *et al*, 2014; Arpin *et al*, 2025). However, the structure of the 50S ribosome-EngA•GDP bound state remains to be determined.

To elucidate the mechanism of EngA-assisted 50S ribosomal assembly, its dissociation subsequent to GTP hydrolysis, and to capture the EngA•GDP ribosome-bound state, we determined the cryo-EM structures of the *Mycobacterium smegmatis (M. smegmatis)* 50S ribosome alone and in complex with EngA•GTPγS or EngA•GDP. Our structural data reveal that both the 50S subunit and EngA undergo substantial coordinated conformational changes during the assembly process, wherein the EngA•GDP states adopt an assembly intermediate conformation in which a stem-loop junction (SLJ) motif is formed between the stem loop of rRNA H80 and the junction connecting H89 and H90. Moreover, ribosome-bound EngA•GDP exhibits a domain conformation that is drastically different from its unbound state.

## Results

### EngA, a 50S ribosome-associated GTPase, is crucial for ribosome biogenesis and mycobacterial growth

In a previous study, we showed that mycobacterial EngA associates with heterologous 50S ribosomal subunit isolated from *E. coli*, an interaction that modulates its enzymatic activity (Agarwal *et al*, 2012). To further assess the intrinsic role of EngA in ribosomal assembly under native conditions, we examined the interaction between *M. smegmatis* EngA, recombinantly expressed and purified from *E. coli* (Supplementary Figure 1A-B), and homologous 50S ribosomal subunits isolated from *M. smegmatis* using sucrose density gradient centrifugation (SDGC) (Supplementary Figure 1C-D). The 50S ribosome-EngA complex formation was subsequently proved by SDGC in the presence or absence of the nucleotide cofactors, including GDP, GTP, and the non-hydrolyzable analog GTPγS. The MsEngA co-sedimented with the mycobacterial 50S ribosomal subunit across all conditions, thus confirming their physical interaction (Figure 1A). Furthermore, the GTPase activity of MsEngA was maintained in the presence of 50S subunits (Figure 1B). Together, these results confirm that the recombinantly expressed EngA is catalytically active and forms a complex with the mycobacterial 50S subunit.

**Figure. 1.**
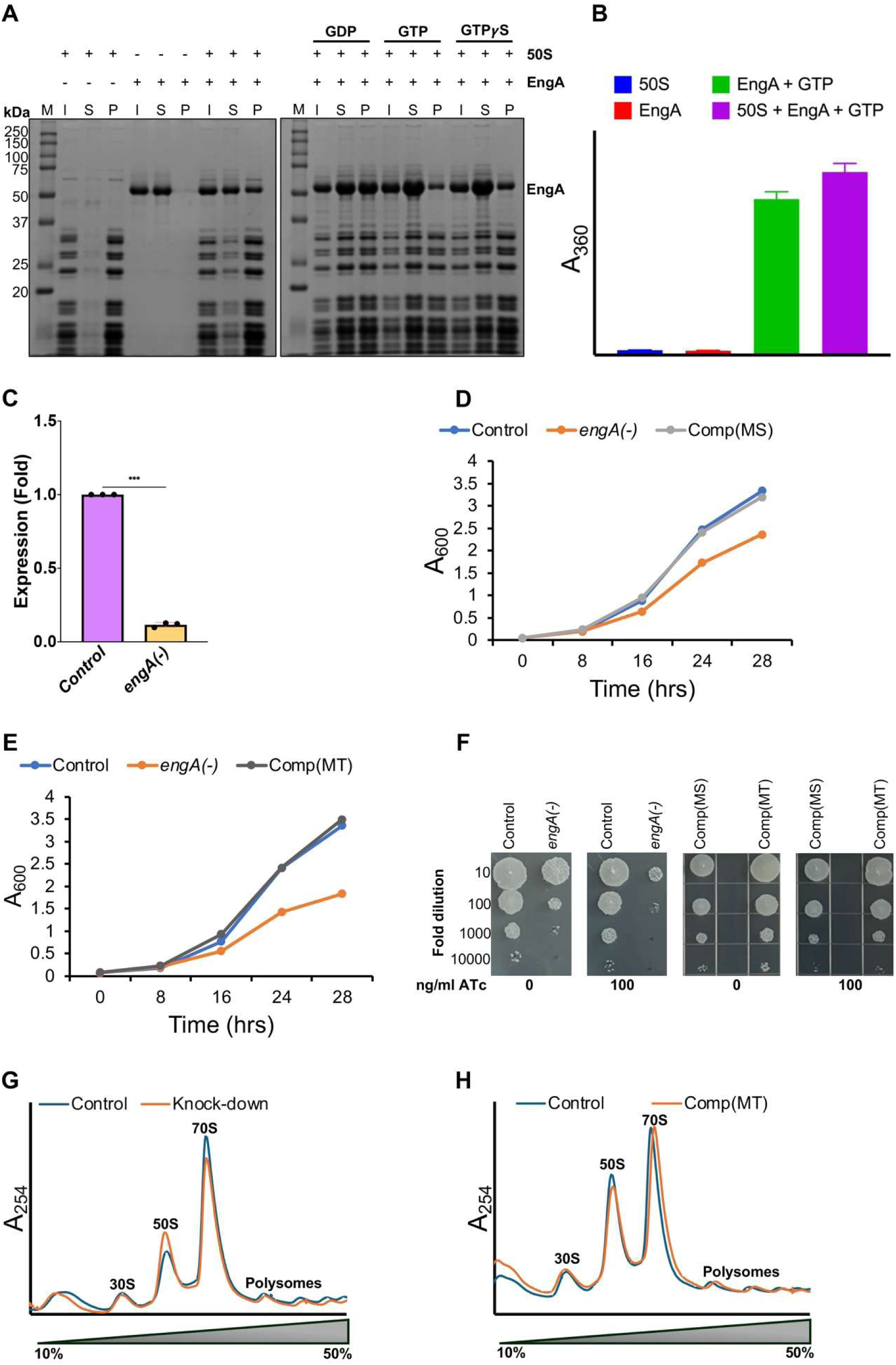
EngA, a 50S-associated GTPase, is essential for *in vitro* growth of mycobacteria. (A) Sucrose pelleting of 50S ribosomes-EngA complex with (+) and without (-) nucleotides (GDP, GTP or GTPγS), was analyzed on 12% SDS-PAGE with Coomassie blue staining. M (marker), I (input), SN (supernatant) and P (pellet) fraction after pelleting on a sucrose cushion. (B) Effect of 50S ribosome on GTP hydrolysis of the wild-type EngA. (C) Expression analysis of *MSMEG_3738* in the *engA(-)* strain of *M. smegmatis.* Expression of *MSMEG_3738* transcripts was estimated by qRT-PCR after 16 h of treatment with 100ng/ml ATc. Shown is the fold expression of *engA* following normalization to the expression levels of the constitutive control gene, *mysA* in the *engA* knockdown strain compared to its level in the empty vector control. (D) Growth curve of Control, *engA(-)* (Knockdown) and *engA(-)::MSMEG_3738* [Comp(MS)] over the time course of 28 hours. (E) Growth curve of Control, *engA(-)* (Knockdown) and *engA(-)::Rv1713* [Comp(MT)] over the time course of 28 hours. (F) Growth phenotype of control, *engA(-)*, Comp (MS) and Comp (MT) was 10-fold serially diluted from 10^−1^ to 10^−5^ analyzed on 7H11 agar media in the presence (100 ng/ml) or absence (0 ng/ml) of ATc. (G) The 10-50% sucrose density gradient fractionation profile of Control and Knockdown; 30S, 50S and 70S are labelled for ribosomal small subunit, large subunit, and mature ribosome, respectively and polysomes. (H)The 10-50% sucrose density gradient fractionation profile of Control and Comp (MT); 30S, 50S and 70S are labelled for ribosomal small subunit, large subunit, and mature ribosome, respectively and polysomes. Data represent mean+SD from three biological replicates in (B-C). Statistical significance in (C) was determined by calculating *p* values with the help of graphpad prism. ***; *p* <0.0005.

To assess the functional implications of EngA-regulated ribosomal assembly in mycobacterial growth, we analyzed the in vitro growth of the *engA* knockdown strain [annotated as *engA(-)*] of *M. smegmatis*. Conditional silencing of *engA* was performed by anhydrotetracycline-induced CRISPRi approach, as described earlier (Choudhary *et al*, 2015). A strain containing the empty vector lacking the *engA*-specific guide RNA served as a control. Knockdown of *engA* transcripts in the *engA(-)* strain was confirmed by quantitative RT-PCR, which revealed ∼90% reduction in the *engA* transcripts following 16 h of treatment with 100 ng/ml ATc (Figure 1C). Growth analysis at regular intervals by estimation of the culture density at 600 nm (OD_600_) revealed that EngA is crucial for sustaining mycobacterial growth under in vitro culture conditions. Knockdown of *engA* severely compromised bacterial growth relative to the control strain (Figure 1D), a phenotype consistent with the defect observed upon its depletion in *E. coli* (Hwang & Inouye, 2001). Importantly, normal growth of *engA(-)* was completely restored upon exogenous expression of either *M. smegmatis engA* or its *M. tuberculosis* orthologue, *Rv1713*. The restoration of growth was confirmed by OD_600_ estimation of the cultures (Figure 1D-E), as well as by evaluating growth of the serially-diluted bacterial cultures spotted on the agar plates in the absence or presence of 100 ng/ml ATc (Figure 1F). Subsequent analysis of the ribosome profiles from the control and the *engA(-)* strains via SDGC revealed an accumulation of 50S subunit peak and a concomitant reduction in the 70S peak relative to the control strain (Figure 1G). These findings imply that upon *engA* knockdown, a fraction of the 50S subunits is trapped in an intermediate assembly state unable to associate with the 30S subunit, leading to a depleted pool of functional 70S ribosomes. As previously observed with the growth, genetic complementation with *M. tuberculosis engA* gene fully restored the ribosome profile, showing a nearly identical proportion of ribosomes and their subunits to that of the control (Figure 1H). Together, these results demonstrate the functional conservation of EngA across mycobacterial species as well as its vital importance in mycobacterial growth and ribosome biogenesis.

### Analysis of cryo-EM structures of the 50S ribosome-EngA complex

To elucidate the molecular mechanism by which EngA assists ribosome assembly, the high-resolution cryo-EM structures of the 50S ribosome in complex with EngA•GTP*γ*S, EngA without added cofactor (natively copurified cofactor), and EngA•GDP were determined (Figure 2, Supplementary Figure 2-5, Supplementary Table 1). For comparison, cryo-EM structures of the 50S ribosome without EngA in pre-mature and fully mature states were also determined (Supplementary Figure 6-7, Supplementary Table 1).

**Figure 2.**
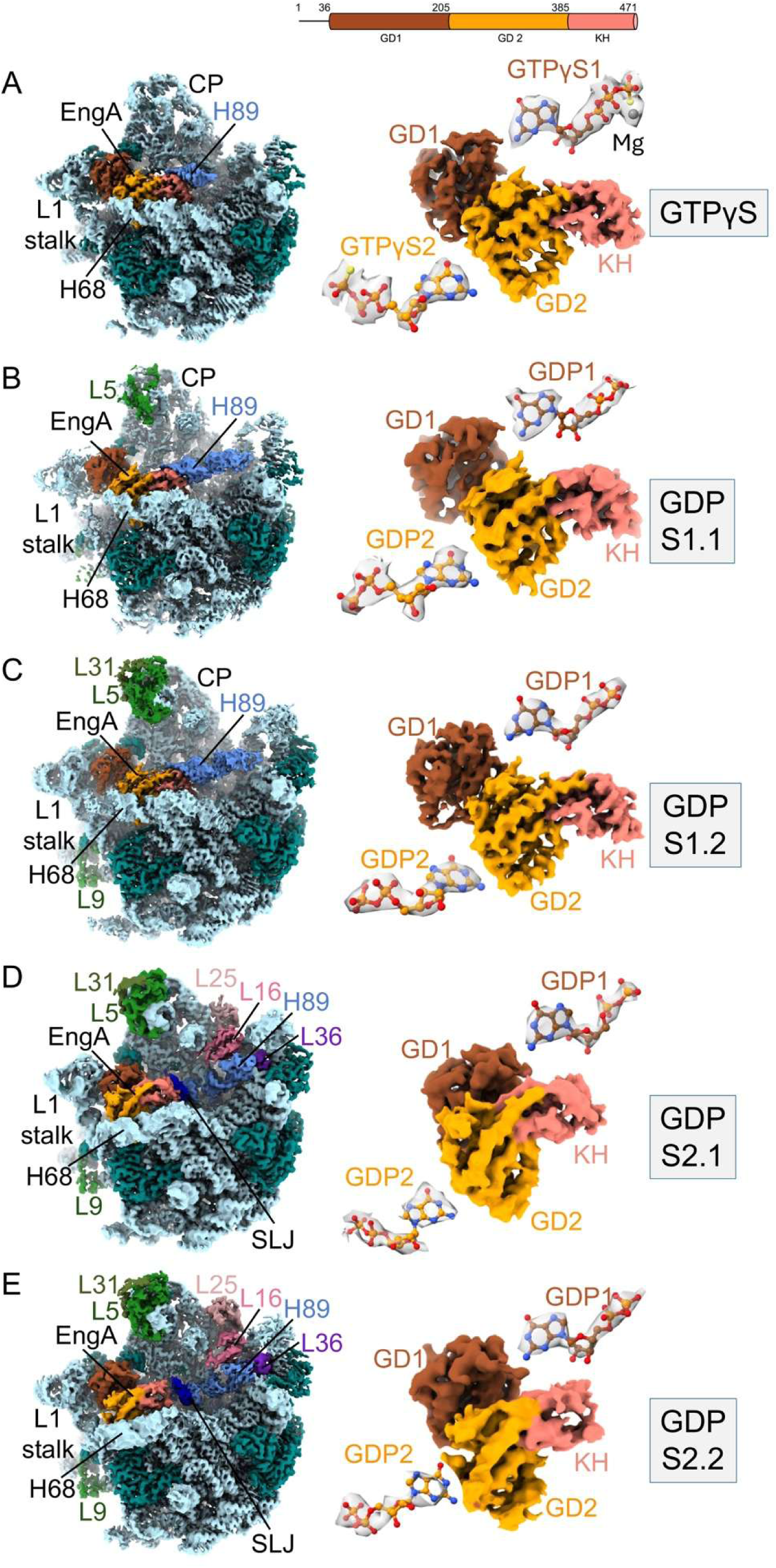
Cryo-EM structures of *M. smegmatis* 50S ribosome in complex with EngA•GTPγS and EngA•GDP. The final cryo- EM maps of the 50S ribosome EngA complex. Left panel, cryo- EM map of 50S ribosome EngA complex, and right panel corresponding cryo- EM densities of EngA and for both nucleotides are shown. (A) 50S ribosome EngA•GTP*γ*S complex, (B) 50S ribosome EngA•GDP (S1.1) complex, (C) 50S ribosome EngA•GDP (S1.2) complex, (D) 50S ribosome EngA•GDP (S2.1) complex, (B) 50S ribosome EngA•GDP (S2.2) complex, are shown. The 23S rRNA (light blue), r-proteins (teal), EngA GD1 domain (sienna), EngA GD2 domain (dark orange), EngA KH domain (salmon), H89 (2420-2719) (cornflower blue), H80 (2470-2482) (medium blue), SLJ (stem loop junction), bL31 (dark olive green), uL5 (forest green), bL9 (geen), bL25 (rosy brown), uL16 (pale violet red), and bL36 (blue violet) are labeled.

#### Structure of the 50S ribosome-EngA•GTPγS complex

To determine cryo- EM structure of the EngA•GTP bound state of 50S ribosome, a non-hydrolyzable analog, GTPγS, was used for in vitro reconstitution of the 50S ribosome-EngA complex. The single-particle reconstruction of the 50S ribosome EngA•GTPγS complex yielded a 2.9 Å-resolution consensus map from the final 312,230 particles after particle polishing in RELION 4.0.0. Further 3D classification into five classes without alignment, classified the particles into four classes with ribosome and EngA complex in different conformations, at 3.1 to 3.3 Å resolution (Supplementary Figure 2A). However, map validation revealed poor map quality and lower surface resolution, driven by a severe preferred particle orientation across all four classes (Supplementary Figure 2B). The preferred orientations were highly likely due to the thin vitreous ice in the grid holes selected for data collection, which has been reported to induce orientation bias (Liu & Wang, 2023).

To overcome the preferred orientation problem, two new cryo- EM datasets of the 50S ribosome EngA•GTPγS complex were collected at an appropriate ice thickness. Single-particle reconstruction yielded a 2.8 Å-resolution consensus cryo- EM map after merging the final selected 274,363 and 321,236 particles from datasets 1 and 2, respectively. Further, structural heterogeneity was sorted using 3D classification without alignment into 10 classes, resulting in four high-quality classes of the EngA-bound 50S ribosome with 272,928 (45.7%) particles (Supplementary Figure 3A). Partial signal subtraction further resolved EngA compositional heterogeneity and yielded a high-quality class containing 182,740 (65.8 %) particles at an overall resolution of 2.8 Å (Figure 2A). The map validation: FSC, local resolution distribution, and particle distribution suggested that the core is resolved up to 2.5 Å resolution, whereas surface resolution is 5.5Å with even distribution of particles (Supplementary Figure 3B, Supplementary Table 1). We could clearly see cryo- EM density for GTPγS in the nucleotide-binding pocket of EngA’s GD1 and GD2 domains, together with the density of the Mg^2+^ ion in the GD1 domain (Figure 2A).

#### Structure of the 50S ribosome-EngA complex without exogenous cofactor addition

To characterize EngA-bound state of the 50S ribosome, an in vitro complex was reconstituted without adding any cofactor. Initial single particle reconstruction of 537,759 particles generated a 3.0 Å consensus map. Further, heterogeneity was sorted using 3D classification and particle polishing. A 3D classification into 10 classes without alignment at T=40, yielding a class with 92,830 (17%) particles having a clear cryo- EM density for bound EngA (Supplementary Figure 4). Cryo- EM map quality was further improved by 3D classification. A major class with a good density of EngA, having 42,175 particles (45.4%), generated a final cryo- EM map with a resolution of 3.4 Å (Figure 2B). The final cryo-EM map was further validated and showed average quality with uneven particle distribution (Supplementary Figure 4). After close inspection of the final cryo- EM map, it was observed that the cofactor in the nucleotide-binding site was occupied by natively purified cofactor GDP (Figure 2B). We call this conformation state GDP state 1.1 (GDP S1.1). Only 8% of the total particles selected for 3D classification yielded the final 50S ribosome bound with EngA•GDP, suggesting that EngA, in its native form, binds with low affinity to the 50S subunit.

#### Structure of the 50S ribosome-EngA•GDP complex

To increase the GDP occupancy and improve the overall quality of the 50S ribosome EngA cryo- EM maps, 50S ribosome EngA complex was in vitro reconstituted with 5,000-fold excess of cofactor GDP. A single-particle reconstruction of the finally selected 934,469 particles generated a 3.4 Å resolution cryo- EM consensus map after particle polishing. Further 3D classification into 10 classes without alignment yielded two classes with better density for bound EngA. Class 6 with 129,113 (13.8%) particles and class 8 with 190,000 (20.3%) particles yielded cryo- EM maps of 3.7 Å and 3.4 Å resolution, respectively (Supplementary Figure 5A). Further classification of class 6 yielded a major class of 49,443 particles (38.3 %), with a better-quality map at 3.3 Å resolution. This class adopted a conformation similar to the GDP S1.1 state; therefore, we named it GDP state 1.2 (GDP S1.2), with clear density for GDP in both nucleotide-binding pockets (Figure 2C). Class 8 particles yielded two distinct EngA•GDP-bound conformations that differ from GDP S1.1 and showed slightly different conformations from each other; therefore, we called these EngA•GDP states 2.1 (GDP S2.1) and EngA•GDP states 2.2 (GDP S2.2) (Supplementary Figure 5B). GDP S2.1, containing 126,761 particles, yielded a final cryo- EM map at 3.1 Å resolution (Figure 2D), whereas the state GDP 2.2, containing 60,523 particles, yielded a final cryo- EM map at 3.3 Å resolution (Figure 2E). All three GDP-bound conformations show high map quality and uniform particle distribution with local resolution up to 2.5 Å (Supplementary Figure 5C). GDP density was well resolved in the GD1 domain of both GDP S2.1 and GDP S2.2 states (Figure 2D-E), whereas the fragmented density in the GD2 domain of GDP S2.2 suggests weaker GDP affinity than in GDP S2.1.

#### Structure of the 50S ribosome in the absence of EngA

To compare the 50S ribosome EngA cofactor-bound structure with that of the fully mature 50S, a cryo- EM reconstruction for the 50S ribosome purified after 70S dissociation through SDGC without EngA was carried out. The total of 169,483 finally selected particles yielded a 3D consensus map at 3.4 Å resolution after particle polishing. Further 3D classification into 5 classes without alignment yielded a mature 50S class with 72,697 (42.8%) particles at 3 Å resolution (Supplementary Figure S6 & S7). A small population of 51,037(6.6%) particles yielded a pre-mature 50S ribosome state (Supplementary Figure S6 & S7). Local resolution suggests that the CP is disordered in the pre-mature 50S ribosome (Supplementary Figure S6B, Supplementary Table S1).

A total of 7 cryo-EM maps were selected for structural analysis. Out of which 5 cryo- EM maps of 50S ribosome with EngA: EngA•GTPγS (GTPγS), EngA•GDP (S1.1), EngA•GDP (S1.2), EngA•GDP (S2.1), and EngA•GDP (S2.2) (Figure 2), and 2 cryo- EM maps without EngA: 50S mature, and 50S pre-mature were selected (Supplementary Figure S7). The cryo- EM data processing and model building statistics are provided in Supplementary Table S1.

### EngA assisted 50S ribosomal subunit assembly

EngA, which possesses N-terminal GD1 and GD2 and the C-terminal KH domains, binds to the conserved binding pocket near the peptidyl transferase center of 50S subunits (Figure 2 and Supplementary Figure 8 and 9). In the pre-mature 50S state, the core of the subunit is assembled, whereas the functional sites: peptidyl transfer center (PTC), central perturbation (CP), L1 stalk, and L7/L12 stalk remain disordered (Supplementary Figure S7). In the GTPγS state, the functional sites begin to assemble upon binding of EngA-GTPγS. GD1 domain interacts with L1 stalk, H82, H88, r-protein uL1, and bL33A, whereas GD2 domain interacts with H68, H75, H88 and uL2, and the KH domain interacts mainly with H80, H89 and H93 (Figures 2A and Supplementary 8A). These interactions are highly conserved, as observed in previous cryo- EM structures reported for *E. coli* and *B. subtilis* (Arpin *et al*, 2025; Zhang *et al*, 2014). The GDP S1.1 state closely resembles GTPγS state, except that uL5 assembles in the CP and H89 becomes slightly ordered (Figure 2B and Supplementary Figure 8B). In GDP S1.2 state, uL31 incorporates, and the N-terminal domain of uL9 becomes ordered (Figure 2C and Supplementary Figure 8C). In all three states, GTPγS, GDP S1.1, and GDP S1.2, EngA adopts an overall similar conformation. Transitioning to GDP S2.1 state, H89 undergoes a major conformational rearrangement that disrupts its interactions with the EngA KH domain. Interestingly, the stem-loop residues of H80 (2474-2478) and those connecting H89 and H90 (2721-2725) adopt a unique stem-loop junction (SLJ) motif (Figure 2D and Supplementary Figure 8D). This conformational rearrangement creates a scaffold for the assembly of r-proteins, uL16, uL25, and uL36 (Figure 2D). The GDP S2.2 state is similar to that of GDP S2.1, except that the EngA KH domain completely dissociates from rRNA helices H80 and H89 and becomes disordered (Figure 2E and Supplementary Figure 8E). Finally, maturation involves further conformational rearrangement of H89 and loss of SLJ motif interactions. Apart from these, large conformational changes occur around PTC, L1 stalk, and the mycobacterial- specific helix H54a becomes ordered (Supplementary Figure S7).

### Nucleotide-dependent conformational changes in GTPase domain of EngA

Each of the two EngA GDs contains five conserved motifs, namely G1-G5, along with two switch regions, namely switch I and switch II (Supplementary Figure S9). Structural comparison of nucleotide binding in both the GDs domains across different intermediate states reveals that the EngA binding pocket is conserved, with nucleotide making extensive hydrogen-bonding, van der Waals, and electrostatic interactions. During GTP hydrolysis, the nucleotide binding pocket undergoes remodeling, which leads to dissociation of EngA from the ribosome (Supplementary Figure 8).

In the GD1 domain, the guanine base of the nucleotide is stabilized through hydrogen- bonding with residues D154, S177, and M179 of the G4 and G5 motifs. The side chain of M179 makes a van der Waals interaction with the nitrogenous base of GTP*γ*S. The ribose sugar moiety interacts with the backbone of Q64. Whereas α and β phosphates interact with Walker-A (G1) motif residues K48, S49, and T50. In the GTPγS-bound state, the γ-phosphate interacts with T70 in the G2 motif (Switch I) (Supplementary Figure 8A middle panel). Most of the nucleotide binding interactions are retained in all the intermediate states, viz., GTPγS, GDP S1.1, GDP S1.2, GDP S2.1, and GDP S2.2 (Supplementary Figure 8, middle panel). However, loss of T70 and γ- phosphate interaction of GTPγS state induces a conformational change in the Switch I region of the GD1 domain. This remodeling results in the formation of hydrogen bonds between the α- phosphate and the main chain nitrogen of Q64, as well as between the side chain of D65 with the 2’-hydroxyl group of ribose sugar in GDP S1.1 and GDP S1.2 states (Supplementary Figure 8B-C middle panel). Further, progression to GDP S2.1 and 2.2 states shows a major conformational change occurring in the G4 motif of the GD1 domain, which disrupts the hydrogen bonds between D154 and the guanine ring (Supplementary Figure 8D-E middle panel). In GDP S2.2, Q64 adopts an altered conformation and loses its interaction with α-phosphate, while Switch I region residues 66-69 become disordered (Supplementary Figure 8E middle panel).

Similarly, nucleotide binding in the GD2 domain involves extensive non-covalent interactions (Supplementary Figure 8 right panel). In GTPγS state, the nitrogenous base interacts with the G4 and G5 motif residues N328, D331, S361, A362, and L363, whereas ribose sugar interacts with K329. The α and β phosphates interact with the G1 motif residues K222, S223, and S224 and γ-phosphate forms a critical hydrogen bond with a highly conserved G2 motif residue T244 (Supplementary Figure 8A right panel). In GDP S1.1 and GDP S1.2, loss of T244 and γ-phosphate interaction is accompanied by disordering of Switch I region residues 230-246. The β phosphate additionally interacts with V220, while the remaining interactions are retained (Supplementary Figure 8B-C, right panel). Remarkably, progression to GDPS2.1 a novel intermediate interaction is established wherein residue K270 re-orients to form a new hydrogen-bonding contact with the β-phosphate group in GDP state 2.1 (Supplementary Figure 8D right panel). However, subsequent progression from GDP S2.1 to GDP S2.2, this interaction with K270 is disrupted as the Switch II region (265-281) becomes disordered, inducing further conformational rearrangement that disrupts the interactions with residues K329, S361, and A362 (Supplementary Figure 8E right panel). Thus, most GDP stabilizing interactions present in earlier states are lost in GDP S2.2, consistent with the fragmented cryo-EM density for GDP in GD2 nucleotide-binding pocket of GDP S2.2. These conformational remodeling of switch regions I and II in both GD domains to become disordered upon GDP bound state are consistent with the crystal structures of EngA homologues, BsYphC•GDP (Muench *et al*, 2006) and TmDer•GDP (Robinson *et al*, 2002).

### Structural rearrangement of EngA domains

EngA, undergoes a huge domain rearrangement upon GTP hydrolysis (Tomar *et al*, 2009). The nucleotide-dependent rearrangement in the binding pocket also induces conformational changes within the two EngA GDs. To understand the domain rearrangement, we superimposed the GD2 and KH domains of GDP-bound states (S1.1, S1.2, S2.1, and S2.2) onto the corresponding domains of EngA in the GTP*γ*S state. Further, we have superimposed the crystal structure of its homologs, BsYphC (PDB: 2HJG), in which GDP occupies both the GDs (Muench *et al*, 2006), and TmDer (PDB: 1MKY), where GDP is bound in the GD2 domain only, and two phosphates are bound in the GD1 domain (Robinson *et al*, 2002). Structural comparison demonstrated that GD2 and KH domains attain a rigid conformation across all the states, whereas the GD1 domain undergoes major conformational rearrangements (Figure 3A).

**Figure 3.**
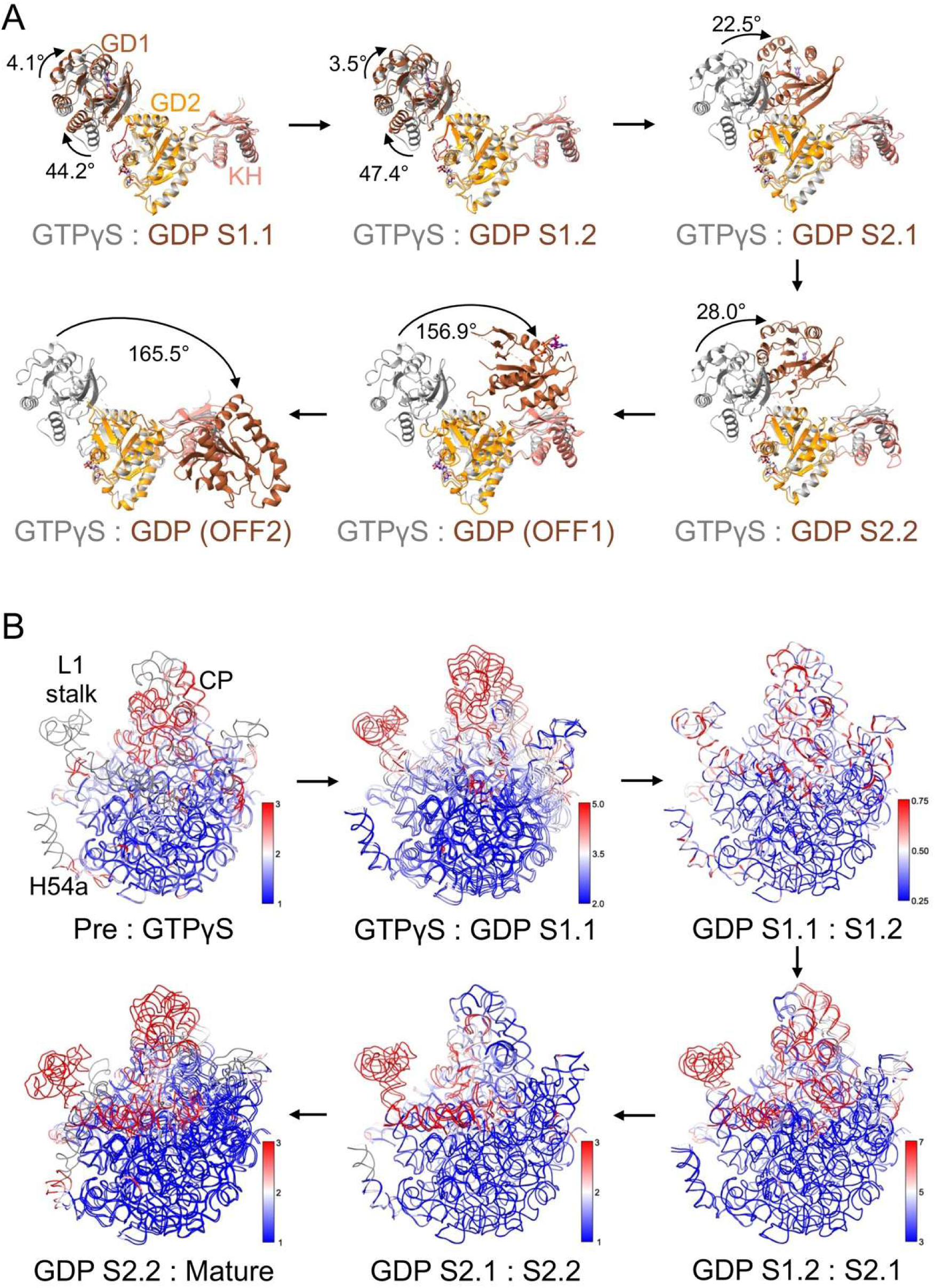
EngA domain conformational changes and associated 23S rRNA remodeling. (A) Structural comparison of domain rearrangements during different nucleotide transition states. The GD2 (dark orange) and KH (salmon) domains in different states: GDP S1.1, GDP S1.2, GDP S2.1, GDP S2.2, and GDP-bound crystal structures OFF1 (PDB ID: 2HJG) and OFF2 (PDB ID: 1MKY) are superimposed to the corresponding domains in the GTPγS (grey) state, and the relative orientation of the GD1(sienna) domain in different states are shown. The arrow and angle (in degree) show the relative position of the GD1 domain in different states w.r.t. its position in GTPγS (grey) state. (B) 23S rRNA superimposed between two states; premature : GTPγS, GTPγS : GDP S1.1, GDP S1.1: GDP S1.2, GDP S1.2 : GDP S2.2, GDP S2.2 : Mature state. rRNA is colored based on r.m.s.d. between corresponding nucleotides, low r.m.s.d. (blue) and high r.m.s.d. (red). A color key is also shown. Nucleotides whose corresponding residues are disordered are shown in grey color.

Subsequent GTP hydrolysis to GDP triggers a conformational change in EngA. The loss of γ-phosphate induces substantial structural rearrangement within α3 helix/switch II region residues (92-114) of GD1 domain and α3 helix swings ∼44.2° and ∼47.4° in S1.1 and S1.2, respectively. These structural changes are accompanied by remodeling of switch I and disordering of GD2 G2 motif residues (236-246), which leads to a rotation of∼4.1° and ∼3.5°, of the GD1 domain in S1.1 and S1.2, respectively (Figure 3A). Next, progression to GDP state 2 (S2.1 and S2.2) is marked by extensive conformational rearrangement in GD1. Relative to the GTPγS state, GD1 undergoes a rotation with respect to GD2-KH domains of about ∼22.5° in GDP S2.1, and ∼28° in the GDP S2.2 state. These structural changes are also accompanied by the GD1 switch I region residues (66-69) becoming disordered (Figure 3A).

Furthermore, structural comparisons of *M. smegmatis* EngA with crystal structures of its counterparts from *B. subtilis* and *T. maritima* showed GD1 domain rotates 156.9° and 165.5° relative to GTP*γ*S, respectively. In these states, GD1 interacts with KH domains, forming a closed state. We proposed these states as closed (OFF) states, BsYphC (PDB: 2HJG) as OFF1 and TmDer (PDB: 1MKY) as OFF2 states, based on the its GD1 rotation angle w.r.t. GTP*γ*S (Figure 3A).

### EngA mediated conformational changes in 23S rRNA

The conformational rearrangement in the domain architecture of EngA is accompanied by dynamic changes in functional center of 23S rRNA (Figure 3B). The CP, which remains disordered in the premature state, becomes partially assembled upon EngA•GTPγS binding and undergoes further assembly in the GDP S1.1 and GDP S1.2 states. The most prominent conformational changes occur during the transition from GDP S2.2 to the mature 50S state, in which CP rotates ∼15° towards PTC (Supplementary Figure 10). The CP also undergoes major displacement up to ∼5Å during the transition from premature to mature 50S state (Figure 3B).

With the progressive reorientation of CP, EngA binding at the PTC region also prevents premature docking of helices H68–H71 and displaces them from their functional positions. H68-71 remains disordered in the premature state (Supplementary Figure 7) while it adopts a similar conformation in GTPγS and GDP S1 (Supplementary Figure 10). However, transition from GDP S1 to GDP S2 induces primary displacement of about ∼22° rotation, translation of ∼4Å (Supplementary Figure 10) and deviation ∼3-7Å (Figure 3B). Following EngA dissociation, these helices undergo a secondary conformational rearrangement, H68 rotating by ∼46°, and H69 translating by ∼14Å towards the PTC (Supplementary Figure 10). These structural rearrangements of H68-H71 yield the fully mature and active PTC, which is capable of associating with the 30S subunit to form a functional 70S ribosome. Overall, H68-H71 helices undergo deviation of about ∼3Å during the process of ribosomal assembly to attain their mature conformation (Figure 3B).

In parallel with the above changes, L1 stalk also undergoes extensive nucleotide-dependent conformation rearrangement. In a pre-mature state, the L1 stalk remains disordered (Supplementary Figure 7) but becomes ordered upon EngA binding through its interactions with the GD1 domain (Figure 2A). Further, GTP hydrolysis and local conformational changes in GD1 domain of EngA cause L1 stalk to deviate up to ∼5Å from its position in GTPγS to GDP S1.1 state (Figure 3B). Its conformation in GDP S1.2 remains similar to GDP S1.1, but it subsequently rotates by ∼16°-18° towards CP in GDP S2.1 and S2.2 states (Supplementary Figure 10). It also undergoes dynamic flexibility of about ∼7Å while transitioning from GDP S1.2 to GDP S2.1 and further dynamic changes of about ∼3Å from GDP S2.1 to GDP S2.2 state (Figure 3B). Finally, EngA dissociation leads to a rearrangement of ∼22° rotation (Supplementary Figure 10) and ∼3Å displacement (Figure 3B), thereby orienting L1 stalk into its mature conformation required for docking of E-site t-RNA.

Along with the maturation of CP, H68-71 and L1 stalk, H89 also undergoes a distinct conformational rearrangement that is accompanied by PTC maturation. In GTPγS, H89 interacts with EngA KH domain (Figure 2A) and retains similar conformation through GDP S1.1 and GDP S1.2 (Supplementary Figure 10). During the transition from GDP S1 to GDP S2.1, H89 undergoes an initial displacement of ∼20Å (Supplementary Figure 10) and forms a transient SLJ motif with residues in apical loop of H80 (Figure 2D-E). This also induces a dynamic flexibility of ∼7Å as H89 transitions from GDP S1.2 to GDP S2.1 (Figure 3B). Subsequently, upon EngA dissociation, H89 undergoes a final structural change with additional ∼11° rotation to achieve its mature conformation (Supplementary Figure 10).

### The stem loop junction rRNA motif is formed in the 50S assembly intermediate

During the 50S subunit assembly process, H89 remodeling is accompanied by a conformational change in H80, creating a scaffold for the late-state r-protein binding. EngA remodels rRNA helices near the PTC, particularly H89, which adopts an interesting intermediate state by forming a SLJ motif with H80 (Figure 2D-E, Figure 4A-C and Supplementary Figure S11A-C).

**Figure 4.**
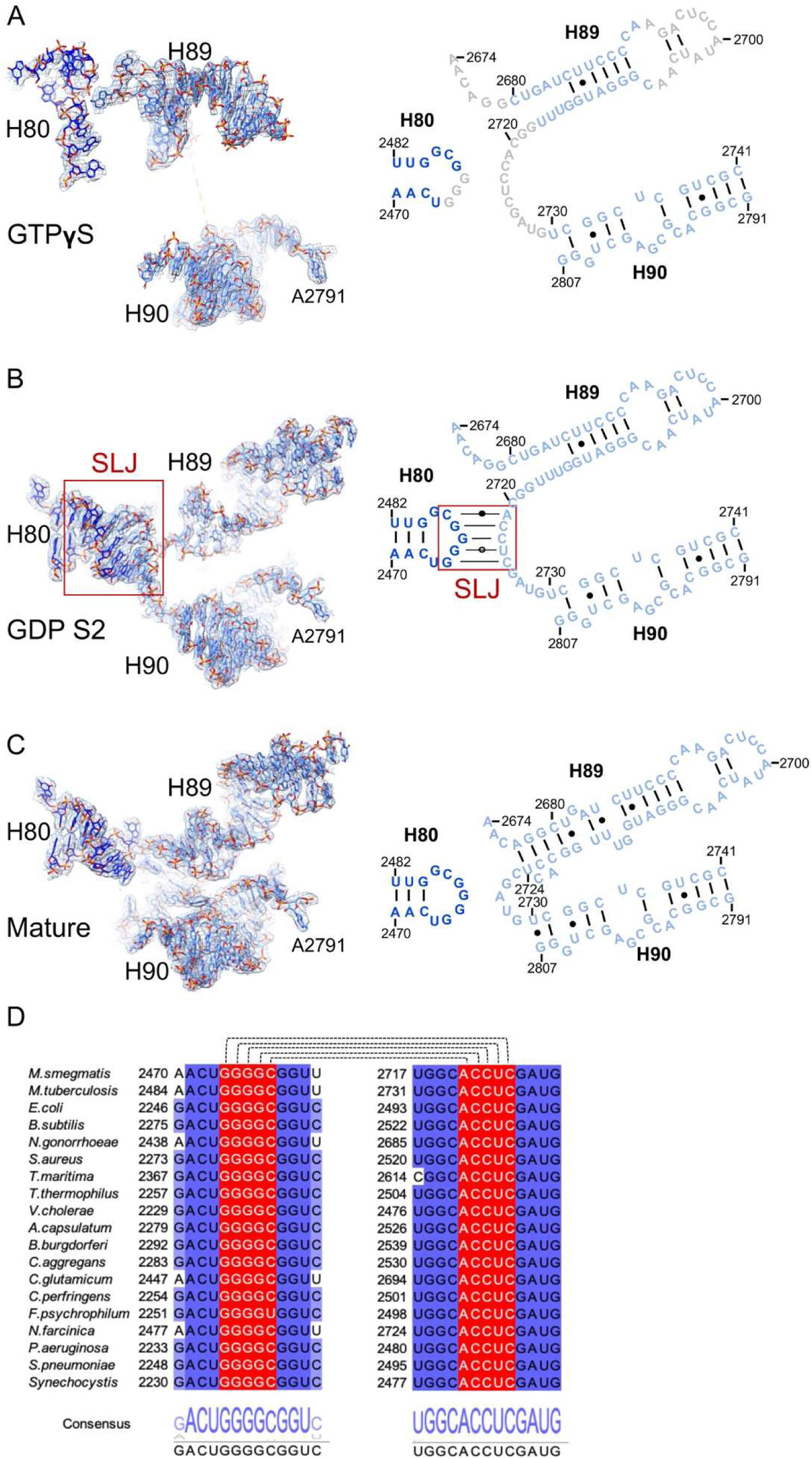
Stem-loop junction interaction. (A) 23S rRNA region encompassing helices H80, H89 and H90 in the GTPγS bound state. The cryo- EM density and corresponding atomic model of the helices are shown on the left. The corresponding 2D representation diagram highlighting the arrangement of helices and their loop nucleotides is shown on the right. H80 nucleotides (medium blue), H89-90 nucleotide (cornflower blue) and disordered nucleotides (grey) are highlighted. (B) 23S rRNA helices in the GDP S2 bound state. The cryo- EM density and corresponding atomic model of helices are shown on the left. The corresponding 2D representation shows the base-pairing interactions formed in KSL during GDP S2 assembly state. (C) Structure of helices H80, H89 and H90 in the mature ribosome is shown. The corresponding cryo- EM density and atomic model of helices are shown on the left. The 2D representation illustrates mature arrangement of helices and associated nucleotides. (D) Multiple sequence alignment of the SLJ regions corresponding to helices H80 and H89 of 23S rRNA from representative bacterial species. Residues involved in the SLJ interaction are highlighted in red, while the rest of the conserved nucleotides are highlighted in dark and lighter purple shades according to sequence conservation, and the consensus sequences are shown below each alignment. Residue and helices are numbered according to the respective 23S rRNA sequences in M. smegmatis. Nucleotide numbering follows the M. smegmatis 23S rRNA sequence. Watson-Crick base pairs are indicated by lines (-), non- canonical base pairs by small dots (●), and nonstandard wobble base pairs by rings (○).

In the GTP*γ*S state, the H89 is partially assembled, and stem loop residues (A2693-A2706) and the residues (G2718-2729) within the segment connecting H89 and H90 remain partially disordered (Figure 4A and Supplementary Figure S11A). The EngA KH domain stabilizes immature H89 conformation through interactions of U2715 phosphate with S387 and F416, U2716 ribose sugar with R385 and A420, and U2717 with with R463. H80 likewise remains unassembled, with disordered loop residues G2474-G2476, while stem region residues A2470-G2474 interact with EngA KH domain. A2471 forms a hydrogen bond with V462, and U2472 interacts with N461 and R463. Essentially, the EngA KH domain stabilizes the premature H80 and H89 conformation (Figure 4A).

While this arrangement is retained in GDP S1.1 and GDP S1.2, in which H80, H89, and the KH domain adopt conformations similar to that in the GTPγS (Figure 2B-C), transition to GDP S2.1 and GDP S2.2 disrupts the interaction between the KH domain with H80 and H89, and KH domain becomes partially disordered (Figure 2D-E). The resulting rearrangements bring the H80 stem loop residues G2474-C2478 into contact with residues A2721-C2725 in the junction connecting H89 and H90, thereby forming a unique stem-loop junction (SLJ) interactions, where three canonical Watson–Crick base pairs (G2474-C2725, G2476-C2723, and G2477-C2722) and one non-canonical wobble base pairs (G2475-U2724) are formed, and two nucleotides, C2478 and A2721 faces each other (Figure 4B). Upon EngA’s dissociation from 50S, the transient stem-loop junction motif is lost, and both H80 and H89 fully assemble. While H80 forms a stem-loop-like structure, the H89 residues of the stem 1 base pairs and the residues of the loop connecting H89 and H90 become ordered (Figure 4C).

### Validation of critical residues by mutagenesis

To understand the functional significance of residues in the nucleotide-binding pocket and at the rRNA interface, site-directed mutagenesis (SDM) of EngA was performed. SDM was used to generate EngA variants targeting the P-loop of the GD1 and GD2 domains (S49A, S223A, and S224A) and the rRNA interface (R157A and R269A), along with the double mutants S49A/S223A and R157A/R269. All the mutant derivatives of EngA were purified using a protocol similar to that used for the wild-type protein, yielding highly homogeneous preparations. Co-pelleting assay was subsequently performed with the individual mutants, which revealed that arginine mutants (R157A, R269A, and R157A/R269A) retained binding affinity for the 50S subunit, whereas the P-loop serine variants (S49A, S223A, S224A, and S49A/S223A) exhibited a slight decrease in binding within the 50S pellet fraction, consistent with the findings from previous studies (Hwang & Inouye, 2010; Bharat *et al*, 2006) (Figure 5A). Impaired 50S binding affinity of the P-loop mutant derivatives is likely driven by the disruption of the hydrogen bonds normally formed between the hydroxyl (-OH) side chains of Ser49, Ser223, and Ser224 and the γ-phosphate moiety of GTPγS (Supplementary Figure 8A, middle and right panels). Moreover, genetic complementation studies with the *engA(-)* strain of *M. smegmatis* using a representative EngA mutant revealed that, unlike the wild-type EngA, the S49A variant was unable to rescue growth on the ATc-containing medium in the spotting assay (Figure 6B), further validating the critical role of the P-loop serine residues in the EngA-50S interaction in vivo.

**Fig. 5.**
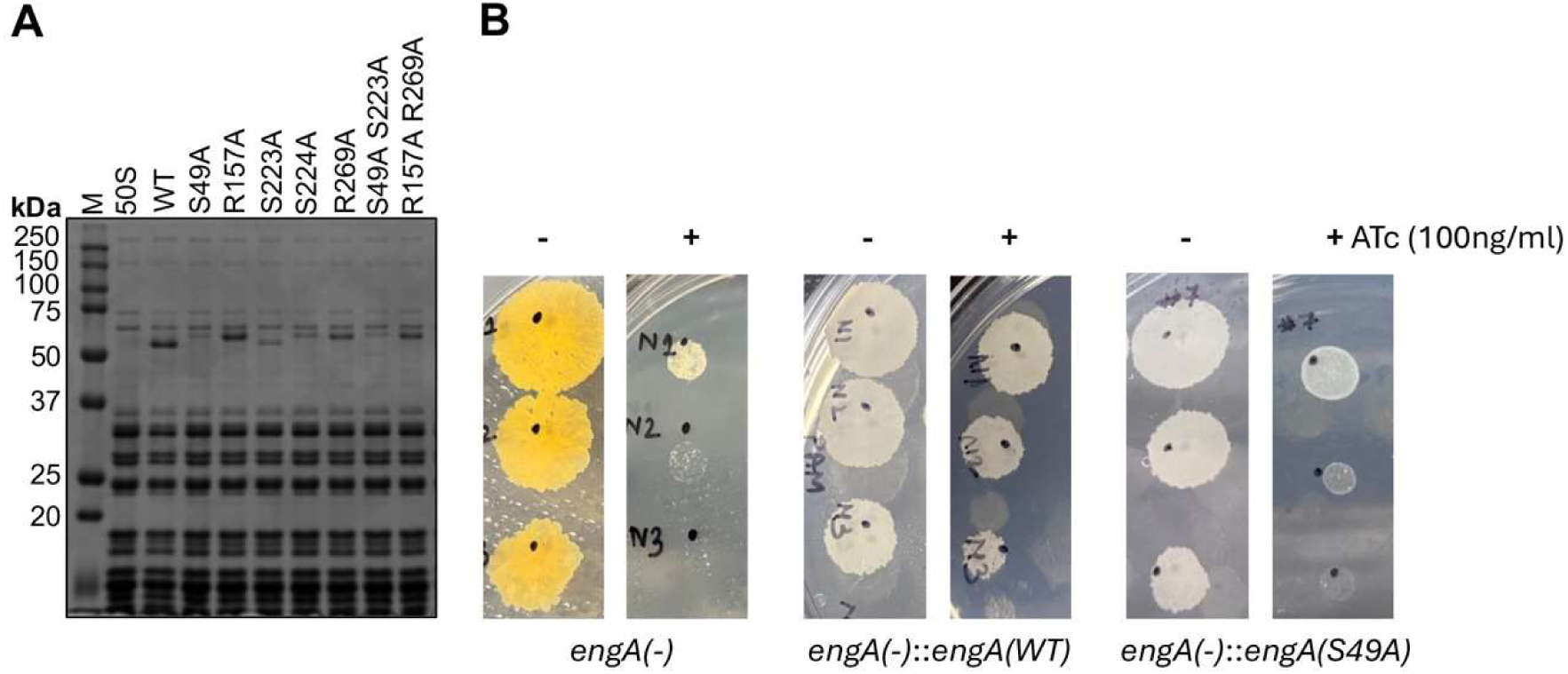
Biochemical and functional characterization of EngA mutants. (A) Interaction analysis of the wild-type (WT) and EngA mutants with 50S ribosome complex. Interaction was determined by sucrose pelleting assay in presence of GTPγS, followed by analysis of pellet fractions on 12% SDS-PAGE with Coomassie blue staining. (B) In vitro growth analysis of the recombinant *M. smegmatis* strains by spotting assay. Serial dilutions of the respective bacterial cultures were spotted on 7H11 agar plates in the absence or presence of 100 ng/ml ATc. Shown are the images of the respective plates after 4 days of incubation at 37 °C.

**Figure 6.**
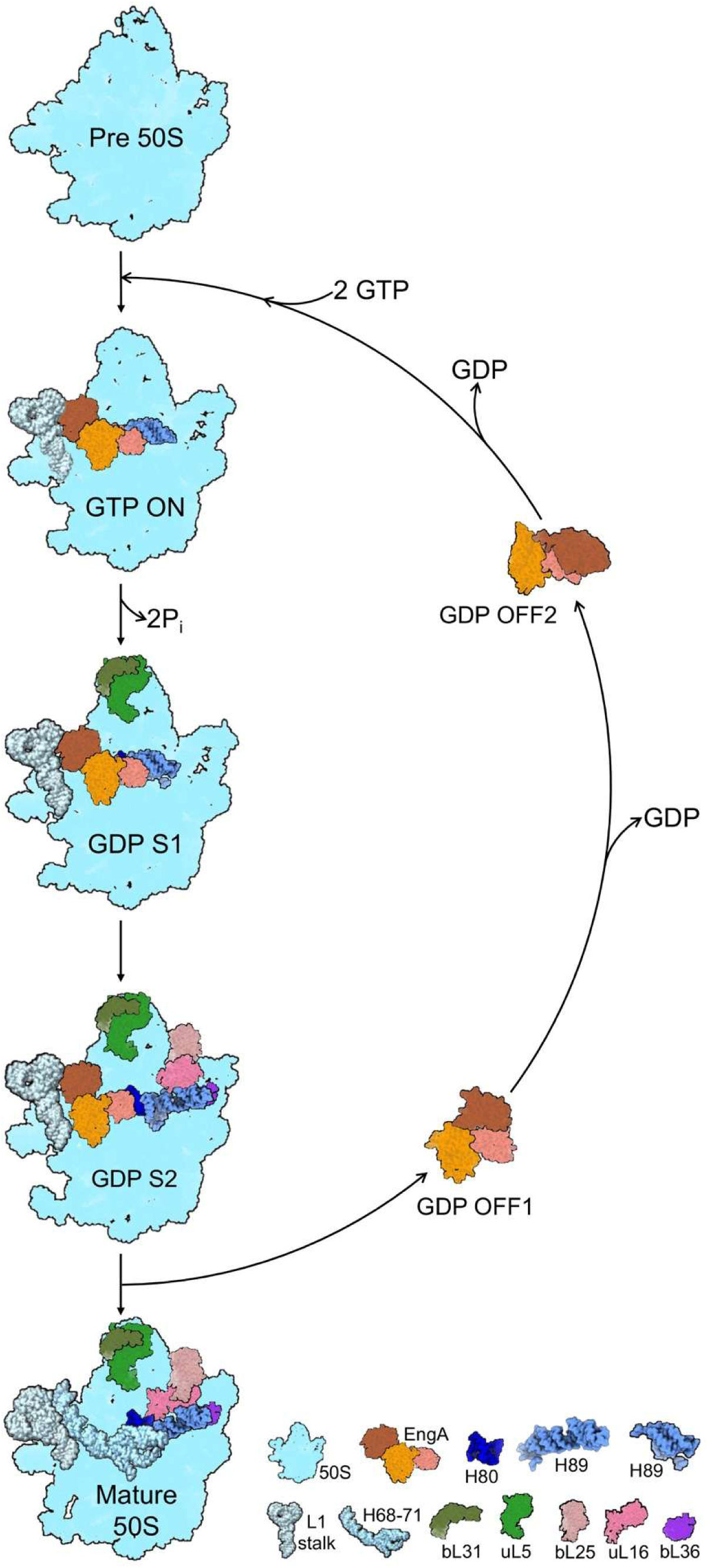
Schematic of EngA-assisted 50S ribosome assembly. Shown is the schematic representation of the EngA-assisted ribosome assembly states– pre-50S, GTP (ON), GDP S1, GDP S2, and mature 50S– reported in this study and in earlier studies with EngA homolog from *T. maritima* (PDB ID: 2HJG) and *B. subtilis* (PDB ID: 1MKY).

## Discussion

### EngA assisted 50S subunit assembly and its impact on mycobacterial growth

In this study, we elucidated the role of EngA, a highly conserved factor involved in late-stage 50S ribosomal subunit assembly, in ribosome biogenesis and its subsequent effects on mycobacterial proliferation. Biochemical experiments demonstrated that EngA binding affinity for the 50S subunit is strengthened in the presence of cofactors, consistent with previous observations (Agarwal *et al*, 2012). Depletion of *engA* led to reduced bacterial growth and accumulation of free 50S subunits, indicating that immature large subunits fail to efficiently associate with the 30S subunit. These defects were rescued by expression of either the *M. smegmatis engA* gene or its *M. tuberculosis* orthologue, demonstrating that EngA’s role is conserved across mycobacteria (Figure 1).

Cryo- EM studies revealed that EngA binds to a conserved binding pocket near the PTC, and that GTP hydrolysis triggers a massive structural rearrangement, specifically the rotation of the GD1 domain toward the CP, eventually docking onto the KH domain, whereas the GD2 and KH domains remain rigid (Figure 2, 3A, and Supplementary Figure 8 and Video 1). Similarly, 23S rRNA also undergoes a structural rearrangement of functionally important sites CP, L1 stalk, H68- H71 and H89 (Figure 2, 3B, and Supplementary Figure 8 and Video 1). During 50S subunit assembly, a stable intermediate state, the GDP S2 state, is formed, which is characterized by the formation of a stem-loop junction motif, and EngA remains bound to the ribosome before exiting from the ribosome (Figure 2 and Figure 4). Mutational analyses further demonstrated that conserved serine residues involved in nucleotide binding are essential for 50S binding and required for cellular viability (Figure 5).

### Stem-loop junction motif is a unique RNA motif in 50S ribosome assembly intermediate

RNA structural motifs are recurring arrangements of nucleotides that play essential roles in diverse biological processes by organizing catalytic sites, regulatory switches, and molecular-recognition interfaces (Batey *et al*, 1999; Moore, 1999). The major secondary-structure motifs, including hairpin loops, internal loops, and bulges, arise from intramolecular nucleotide base pairing, which provides a framework for higher-order RNA folding [32, 34]. The tertiary motifs are formed through interactions between two distinct secondary elements. These help to establish compact and stable three-dimensional structures(Gluick & Draper, 1992). These tertiary motifs include A-minor interactions, ribose zippers, coaxial stacking, tetraloop-receptor interactions, base triples, kissing-stem loop and pseudoknots (Nissen *et al*, 2001).

We identified a unique tertiary motif in 50S ribosomal assembly intermediate GDP S2 state, in which the EngA KH domain dissociates from H89, allowing H89 to undergo a conformational change in such a way that the junction connecting H89 and H90 base pairs with the apical loop of H80, forming a stem-loop junction motif (Figure 2 and 4 & Supplementary Figure 8 and11 and Video 1). This motif creates a scaffold for incorporating r-proteins, L16, L25, and L36. The sequence involved in the formation of this motif is highly conserved among all eubacteria (Figure 4D and Supplementary Figure 11D), and EngA is also conserved across eubacteria (Supplementary Figure 9). Suggesting that stem-loop junction motif formation is a conserved pathway in the EngA-mediated assembly of the 50S ribosome.

### A multi-state model for EngA functioning

EngA is a novel GTPase, as it possesses two consecutive GTPase domains, GD1 and GD2, followed by a KH domain and is involved in late-stage assembly, when the ribosomal core is already assembled (Tomar *et al*, 2009; Zhang *et al*, 2014). Earlier, X-ray crystallography and biophysical studies on EngA proposed that it switches between two states: the open (ON) state and the closed (OFF) state. EngA with GDP bound in both domains as a closed (OFF) state and GDP bound in the GD2 domain, where only two phosphates in the GD1 domain (without GDP) as an open (ON) state. Two phosphates observed in the EngA GD1 domain of the crystallographic structure were assumed to be a mimic of the GTP-bound state. Therefore, it was proposed to be in an active open (ON) state (Muench *et al*, 2006). The cryo- EM structure of EngA•GMPPNP complex with the 50S ribosome revealed that EngA binds near PTC and adopts an open conformation (Zhang *et al*, 2014), which is quite different from the earlier proposed open (ON) conformation of EngA in the ribosome unbound state. The EngA•GDP-bound close (OFF) state was docked into the 50S ribosome EngA•GMPPNP complex to demonstrate the EngA•GDP ribosome-bound state. However, the EngA•GDP bound complex with the 50S ribosomal structure remained elusive.

Our cryo- EM studies provided the 50S ribosome-bound EngA•GDP in multiple states. Consolidating earlier structural data with our findings, we proposed a multi-state EngA model for functioning (Figure 6 and Supplementary Video 1). EngA•GTP recognizes the pre-50S particle, in which only the ribosomal core is assembled and binds in the interface region near the L1 stalk, H68-71, H80 and H89 region, where it recognizes and stabilizes the immature conformation of these elements. We propose this as an active open (ON) state. Hydrolysis of GTP molecules and release of 2Pi acts as a switch that moves the complex to the intermediate GDP S1 state. The EngA GD1 domain rotates ∼3.5° to ∼4.1° w.r.t. and and helix 5 swings between ∼44.2° to ∼47.4°. This allows the first set of ribosomal proteins, bL31 and uL5, to be recruited near the central protuberance, while EngA, with two GDPs bound, remains in the bound state and the L1 stalk attains an intermediate conformation. Assembly then advances to the intermediate GDP S2 state, EngA remains bound in its GDP form while assembly continues. EngA GD1 rotates ∼22.5° and ∼28.0° towards KH domain. This rearrangement of EngA dislodges its KH domain from H89, thereby inducing a conformational change in H89. This enables loop connecting H89-H90 to form a stem-loop junction motif with the apical loop of H80. The proteins bL25, uL16 and bL36 are incorporated. EngA is proposed to stabilize these elements in this conformation until the surrounding regions are mature, thus preventing their premature docking. Subsequently, EngA•GDP form dissociates from the 50S ribosome, and the ribosome takes its mature conformation by remodeling its rRNA helices, L1-stalk, CP, H89, and H68-H7. In the ribosome unbound or free state, EngA with GDP bound in both GD domains undergoes a ∼156.9° rotation and docks on the KH domain. This is the inactive close (OFF) state as proposed previously (Muench *et al*, 2006). EngA undergoes further conformational change upon dissociation of GDP from the GD1 domain, and with GDP bound in the GD2 domain only. Where the GD1 domain rotates by ∼165.5° and docks on the other side of the KH domain. We now propose this state as GDP OFF2, which was previously proposed as an open (ON) state (Muench *et al*, 2006). When EngA senses the higher concentration of GTP in the cell, both the GD domains will be occupied by GTP and EngA will restore its open conformation, which would favor its binding to the ribosome (Figure 6).

Overall, this study establishes a multistep model for EngA-assisted ribosome assembly. Where EngA and rRNA undergo a series of coordinated structural changes that guide the pre-mature 50S ribosome to assemble into its mature form. Further, this study revealed an assembly intermediate, the GDP S2 state, characterized by a unique stem-loop junction (SLJ) motif. Whether these are the only intermediate states or whether there are other transient states that could not be detected by cryo- EM, remains to be demonstrated. A molecular dynamics simulation, supplemented by biochemical studies, may reveal yet another uncharacterized intermediate state.

## Methods

### Bacterial strains and culturing

This study involved *E. coli* XL2 Blue strain (Agilent Technologies) for cloning and propagation of plasmids, whereas the *E. coli* BL21(DE3) (Agilent Technologies) was used for expression and purification of recombinant proteins. *M. smegmatis* mc^2^155 strain was kindly provided by Dr. William Jacobs at the Albert Einstein College of Medicine, USA. *E. coli* was cultured in LB medium (Himedia), whereas *M. smegmatis* was cultured in Middlebrook 7H9 supplemented with 0.05% tyloxapol and 0.5% glycerol (Merck) or in Middlebrook 7H11 without detergent. Bacteria were cultured in broth medium at 37 °C with shaking at 200 rpm, whereas growth on agar plates was obtained in a standing incubator at 37 °C. Wherever required, we used 50 µg/ml kanamycin and 150 µg/ml hygromycin for culturing recombinant *E. coli*, and 25 µg/ml kanamycin and 50 µg/ml hygromycin for mycobacterial transformants.

### Construction of *engA* knockdown and complemented strains

To generate the *engA* knockdown strain of *M. smegmatis*, the *MSMEG_3738*-specific guide sequence cassette was obtained from pGrna-*engA^MS^* clone (Choudhary *et al*, 2015) by restriction digestion with *Nhe* I and *Not* I, and cloned in an integrative Kan^R^ plasmid pTetInt-dCas9 to express both the guide RNA and dCas9 from the same plasmid. The resulting plasmid, annotated as pTetInt-dCas9_MS3738 was electroporated in *M. smegmatis* and colonies were selected on 7H11 agar plates in the presence of kanamycin. The recombinant colonies were verified by PCR before inoculation in the broth medium.

To achieve constitutive expression of *MSMEG_3738* in the *engA* knockdown strain of *M. smegmatis*, a ∼350bp promoter fragment of the *yidC* operon containing SigA-recognition sequence 5’-TTGACC-N17-TACCCT-3’ (Agarwal & Tyagi, 2006) was PCR amplified using the yidC_pro(F) and yidC_pro(R) primers (Supplementary Table 2). This fragment was cloned into the replicative *E. coli*-mycobacteria shuttle plasmid pTetR (Agarwal & Tyagi, 2006) at the *Nhe* I and *Nde* I sites. The resulting HygR plasmid was annotated as pYidCpro. *MSMEG_3738* was PCR- amplified using a two-step overlapping PCR strategy with specific primer pairs, MS3738(F)- PAM(R) and PAM(F)-MS3738(R) (Supplementary Table 2) to introduce a synonymous mutation disrupting its protospacer adjacent motif (PAM) sequence. The *MSMEG_3738(ΔPAM)* fragment was subsequently digested with *Nde* I and *Hind* III and cloned into the corresponding sites of pYidCpro to yield pYidCpro-MS3738, which was electroporated into the *engA(-)* strain of *M. smegmatis*. The resulting Kan^R^-Hyg^R^ strain was subsequently used for complementation studies. To constitutively express *Rv1713* in the *M. smegmatis engA(-)* strain, *Rv1713* was PCR amplified using genomic DNA of *M. tuberculosis* H37Rv as a template with a pair of oligonucleotides- Rv1713F and Rv1713R (Supplementary Table 2), yielding a ∼1.4 kb PCR amplicon carrying *Nde* I and *Hind* III overhangs at the 5′ and 3′ ends, respectively. The *Nde* I-*Hind* III digested PCR- amplified *Rv1713* gene fragment was subsequently cloned into pYidCpro at the same sites. The sequence-verified pYidCpro-Rv1713 clone was used to electroporate into the *engA(-)* strain of *M. smegmatis*.

To analyze the growth of the recombinant *M. smegmatis* strains, cultures were inoculated at an initial OD600 of ∼0.05 with and without 100 ng/ml ATc, and OD600 was measured after every 4 hours. For spotting images, 1:10 serial dilutions of the freshly grown turbid cultures were prepared. Furthermore, 3 µl of each serial dilution was spotted onto agar plates in the presence or absence of 100 ng/ml ATc.

### RNA extraction and Quantitative Real-Time PCR (qRT–PCR)

*M. smegmatis* cells were lysed by bead beating in RNAiso Plus to extract total RNAs, according to the manufacturer’s instructions (Takara Bio). RNAs were quantitated by nanodrop and equal amount of RNAs from the test and the control samples were subjected to DNase I-treatment using RNase-free DNAse I (Ambion) to degrade contaminating genomic DNA. First-strand cDNA was synthesized using 500 ng of DNAse-treated total RNA with the help of SuperScript IV reverse transcriptase and random hexamers (Thermo Fisher). qRT-PCR was subsequently performed on an ABI 7500 Fast System using SYBR Green PCR master mix (Applied Biosystems) and gene-specific primers (Supplementary Table 2). Relative expression, normalized to the housekeeping gene *mysA*, was determined using the comparative CT method (2^−^ΔΔCT_)._

### EngA protein purification

The *MSMEG_3738* gene, which encodes EngA, was cloned into the pET-28b (+) bacterial expression vector (Agarwal *et al*, 2012) and transformed into *E. coli* BL21(DE3) competent cells. Cultures were grown at 37 °C to an OD600 of 0.6, induced with 0.5 mM isopropyl β-D-1-thiogalactopyranoside (IPTG), and incubated for 16 h at 18 °C. Cells were harvested by centrifugation at 6,000 rpm for 30 min at 4 °C and resuspended in lysis buffer containing 50 mM Tris-Cl (pH 8.0), 300 mM NaCl, 5% glycerol, 10 mM imidazole, 1 mM PMSF, 0.5% Tween-20, 5 mM MgCl2, and 5 mM β-mercaptoethanol. After sonication, the lysate was clarified at 13,000 rpm for 1 h at 4 °C. The supernatant was incubated with Ni-NTA His-Bind resin (Millipore) for 3 h at 4 °C with gentle rocking. The resin was washed with 30 ml of buffer containing 50 mM Tris-Cl (pH 8.0), 300 mM NaCl, 5% glycerol, 20 mM imidazole, 2 mM MgCl2, and 5 mM β-mercaptoethanol. EngA was eluted with 20 ml of the same buffer containing 300 mM imidazole. The eluate was concentrated with a 30-kDa-cutoff Amicon filter (Merck) and further purified on a Superdex 200 Increase 10/300 column (Cytiva). Peak fractions were analyzed by 12% SDS-PAGE; pure fractions were pooled, concentrated, and reanalyzed by SDS-PAGE (Supplementary Fig. 1A and B). Protein concentration was determined from absorbance at 280 nm. EngA was concentrated to ∼2.5 mg/ml and stored at −80 °C.

Site-directed mutagenesis was used to generate substitutions in the GD1 and GD2 nucleotide-binding sites of EngA (Supplementary Table 2). Mutants were constructed by overlap-extension PCR. The *engA* gene was first amplified as two fragments from pYidCpro-MS3738 (PAM): one using ORF-F and the relevant mutant reverse primer, and the other using the matching mutant forward primer and ORF-R. The two amplicons were then combined as templates for overlap-extension PCR with ORF-F and ORF-R. The resulting product was gel-purified, digested with *NdeI* and *HindIII*, and cloned into the corresponding sites of pET-28b(+). All recombinant plasmids were verified by DNA sequencing. For double mutants, a plasmid carrying the first substitution served as the template for a second round of mutagenesis with mutation-specific primers. Mutant proteins were purified using the wild-type EngA protocol.

### GTPase assay

The GTPase activity of wild-type (WT) EngA was assessed in 96-well plate using the EnzChek Phosphate Assay Kit (Invitrogen E6646), following the manufacturer’s protocol. The reaction mixtures were prepared by combining reaction buffer, 2-amino-6-mercapto-7-methylpurine riboside (MESG) substrate, purine nucleoside phosphorylase (PNP) enzyme and deionized water. Concentration of MESG and PNP was taken according to the protocol described in kit. EngA WT (5 μM) protein was added to the reaction mixture. Upon addition of 1 mM GTP into the reaction mixture, hydrolysis of GTP was determined by measuring absorbance at 360 nm at an interval of every 5 min. The GTPase activity of EngA WT was also measured in the presence of 50S ribosome (0.5 μM) (Figure 1B). Similarly, the GTPase assay was performed for all the EngA mutants using the same protocol (Figure 6B).

### Ribosome isolation and purification from *Mycobacterium smegmatis*

Ribosomes from *M. smegmatis* (mc²155) were isolated using a previously reported protocol (Kumar *et al*, 2024) with minor modifications. The culture was grown in Sauton’s media at 37 °C till mid-log phase and then harvested by centrifugation at 6,000 rpm for 30 min at 4 °C. Cells were lysed under cryogenic conditions using a Mixture Mill MM500 (Retsch) at 30 Hz for six cycles of 1 min each. The lysate was resuspended in lysis buffer (20 mM HEPES pH 7.4, 20 mM MgCl₂, 100 mM NH₄Cl₂, 1 mM PMSF, 3 mM DTT, 1X Protease Inhibitors Cocktail Tablet) and cell debris was removed by centrifugation at 13,000 rpm for 30 min at 4 °C. The supernatant was gently layered over a 1.1 M sucrose cushion buffer (20 mM HEPES pH 7.4, 20 mM MgCl₂, 100 mM NH₄Cl₂, 3 mM DTT) and subjected to ultracentrifugation at 32,700 rpm (Type70 Ti rotor, Beckmann Coulter) for 16 h at 4 °C. The crude ribosome pellet was resuspended in pellet dissolving buffer (20 mM HEPES, pH 7.4, 20 mM MgCl₂, 30 mM NH₄Cl₂, 3 mM DTT), homogenized by Dounce homogenizer, and treated with DNase І (5 U/μl, ThermoFisher) for 1 h on ice. After centrifugation at 13,000 rpm for 30 min at 4 °C, the ribosomal concentration was determined by absorbance at 260 nm. For further separation of ribosomal subunits, 64 O.D. units of crude ribosomes were loaded onto 10–50% sucrose gradients in pellet dissolving buffer and ultracentrifuged at 22,000 rpm for 10 h using SW28 rotor (Beckmann Coulter). Fractions were collected with a Gilson fractionator on a BioComp system (Supplementary Figure 1C). The 70S and polysomes fractions were pooled and concentrated using 100 kDa cut-off Amicon (Millipore) and stored in storage buffer (20 mM HEPES pH 7.4, 20 mM MgCl₂, 30 mM NH₄Cl₂, 3 mM DTT).

### Ribosome dissociation

To dissociate 70S ribosomes into individual ribosomal subunits, i.e. 30S and 50S ribosomal subunits, MgCl₂ concentration was reduced from 20 mM to 1 mM by buffer exchange with dissociation buffer (20 mM HEPES pH 7.4, 1 mM MgCl₂, 30 mM NH₄Cl₂, 3 mM DTT, 2 mM spermidine), followed by incubation on ice for 3-4 h. Dissociated subunits were separated by ultracentrifugation through 10-50% sucrose gradients in dissociation buffer at 25,000 rpm for 8 h (SW28 rotor, Beckmann Coulter), followed by gradient fractionation and concentrated using 100 kDa Amicon filter. The ribosomal subunit fractions were collected and analyzed on 2% agarose gel electrophoresis containing 0.06% bleach and stained with 0.2 μg/ml ethidium bromide (Supplementary Figure 1D). Peaks corresponding to 50S and 30S ribosomal subunits were pooled and concentrated separately, and their concentrations were quantified by measuring absorbance at 260 nm. The concentrated 50S ribosomal subunit was used for further assays.

### 50S ribosome EngA complex formation

The 50S ribosome EngA complex was formed with and without cofactor, GTP or or GDP, or GTPγS, a non-hydrolyzable GTP analog. The 50S ribosome EngA complex was assembled in a 100 µl reaction by incubating 1 µM 50S ribosomal subunits with 10 µM EngA and incubated at 37 °C for 15 min in complex binding buffer (20 mM HEPES pH 7.4, 4 mM MgCl₂, 100 mM NH₄Cl₂, 2 mM DTT, and 2 units/µl RNase inhibitor), followed by an additional 10-minute incubation on ice. For 50S ribosome EngA•GTPγS complex formation, 2 mM of GTPγS was added to the above reaction mixture. Similarly, the 50S ribosome EngA•GDP complex was formed by adding 2 mM GDP (Figure 1A). Further complex formation with mutants was observed in the presence of GTPγS cofactor (Figure 6A).

### Sucrose pelleting assay

The reaction mixture was then centrifuged at 13,000 rpm for 20 min at 4 °C, and subsequently, 90 µl of the reaction mixture was layered onto 500 µl of 0.8 M sucrose cushion prepared in a 1 ml open thick-walled polypropylene tube. The complex was pelleted by ultracentrifugation at 390,000 × g for 6 hours using Beckmann Coulter MLA-150 rotor (Schubert *et al*, 2020). The pellet was resuspended in 30 µl of complex binding buffer, while the supernatant was concentrated to 30 µl using a 30 kDa Amicon filter (Millipore). The presence of EngA was assessed by analyzing the pellet and supernatant fractions from reactions on a 12% SDS-PAGE gel, followed by Coomassie blue staining (Figure 1A). Further, binding affinity for all mutants was checked with 50S ribosomal subunit and analyzed on 12% SDS-PAGE gel (Figure 5A).

### Initial cryo-EM sample preparation and screening

Initial sample screening using negative staining and cryo- EM condition optimization using cryo- TEM were carried out at the Advanced Technology Platform Centre, Regional Centre for Biotechnology, Faridabad. The negative staining was performed by applying 3 µl of 3 mg/ml 50S ribosome-EngA complex with and without cofactors onto a glow-discharged 300 CF300-Cu grids (Electron Microscopy Sciences). The excess sample was blotted, washed with MilliQ water, and stained with an alternative 2% uranyl acetate solution (Sigma). Grids were examined in a JEOL 1400 JEM microscope operating at 120 kVa. For cryo- EM sample screening, complexes were prepared just before grid preparation. 0.4 µM of 50S ribosomes were mixed with 4 µM EngA with and without 2 mM GTPγS or 2 mM GDP in complex binding buffer and incubated at 37 °C for 15 min. This was followed by incubation on ice for 10 min, and cryo- EM grids were prepared using Gatan CP3 plunger by applying 3 µl of 50S ribosome EngA complex onto a 50 s glow-discharged grid R 1.2/1.3 on 300 mesh Cu Quantifoil (TED Pella, INC), followed by 3 s blotting and plunge- freezing in liquid ethane. Grids were imaged using a JEOL 2200FS JEM microscope operating at 200 kVa equipped with a Gatan K2 Summit direct electron detector. Movie stacks were collected at 1.3 Å pixel size with a total dose of 1.1 e⁻/Å² per frame across 30 frames.

### High resolution cryo- EM data collection

Initial 50S ribosome-EngA•GTPγS complex, high-resolution data were collected using a Titan Krios microscope (ThermoFisher) operated at 300 kVa with a Gatan K3 detector at 1.1 Å pixel size and 81,000 x magnification, at SATHI-National Cryo-EM Facility, IIT Delhi, India. A total of 8930 movie stacks, each containing 35 frames with an electron dose of 1.06 e⁻/Å² per frame, in electron-counted movie mode (Supplementary Table 1) were collected.

The final 50S ribosome-EngA•GTPγS complex datasets were collected using a Titan Krios microscope (ThermoFisher) operated at 300 kVa with a Falcon 3 detector. Data was collected in two sets in electron-counted movie mode. For Data 1, a total of 4812 movie stacks, each containing 18 frames at 1.07 Å pixel size, 75,000 x magnification, with an electron dose of 1.09 e⁻/Å² per frame. For Data 2, 4708 movie stacks were collected at 1.07 Å pixel size with a total dose of 0.99 e⁻/Å² per frame across 32 frames (Supplementary Table 1). These datasets were collected at the National Electron Cryo-Microscopy, Bangalore Life Science Campus, Bangalore, India.

The 50S ribosome EngA•GDP complex high-resolution data was collected with Titan Krios microscope (ThermoFisher) operated at 300 kVa with a Gatan K3 detector at ANRF National Cryo- EM facility at IIT Kanpur, India. Data was collected at a pixel size of 1.1 Å with 81,000 x magnification. 17,632 movie stacks were collected, containing 30 frames, an electron dose of 1.1 e⁻/Å² per frame, in electron-counted movie mode (Supplementary Table 1).

For the 50S ribosome EngA complex, the high-resolution data was collected with Titan Krios microscope (ThermoFisher) operated at 300 kVa with a Gatan K3 detector at ANRF National Cryo- EM facility at IIT Kanpur, India. Data was collected at a pixel size of 1.1 Å with 9443 movie stacks, each containing 30 frames, an electron dose of 1.1 e⁻/Å² per frame, in electron- counted movie mode (Supplementary Table 1).

The 50S ribosome, the high-resolution data was collected using a Titan Krios microscope (ThermoFisher) operated at 300 kVa with a Falcon 4i detector at the ANRF National Cryo- EM facility at IIT Madras. A total of 4,034 movie stacks were collected in EER format at a pixel size of 1.22 Å, a magnification of 105,000×, and a total electron dose of 40 e⁻/Å² in electron-counting mode (Supplementary Table 1).

### Single particle reconstruction

RELION 3.1.4, RELION 4.0.0 and RELION 5.0.0 (Scheres, 2012) were used for single particle data processing of all four datasets, 50S-EngA•GTPγS, 50S-EngA•GDP, 50S-EngA, and 50S ribosome (Supplementary Figures 2-6 and Supplementary Table 1). The data were collected as movie stacks, which were aligned and corrected for beam-induced motion using RELION’s own implementation to generate single micrographs. These aligned and motion-corrected micrographs were then CTF-corrected using CTFFIND-4.1.14 (Rohou & Grigorieff, 2015). Further, particles were picked manually from these aligned images and were used as 2D reference for automatic particle picking. Auto-picked particles were then subjected to multiple rounds of 2D classification to remove impurities and degraded particles from the data. The best 2D class averages were selected for 3D classification. A 60 Å low-pass filtered 50S ribosome cryo-EM map (EMDB ID 37552) from *Mycobacterium smegmatis* (Kumar *et al*, 2024) was used as a reference map. Multiple rounds of 3D classification were performed to remove low-resolution 3D classes. The best 3D classes containing 50S ribosomes with EngA bound, and only 50S ribosomes in the control data, were then further refined by estimating asymmetrical and symmetrical aberrations, as well as anisotropic magnification, using CTF refinement (Zivanov *et al*, 2020). Further, these particles were subjected to reference-based beam-induced motion correction in Bayesian Polishing (Zivanov *et al*, 2019). To improve the density of EngA, partial signal subtraction (Bai *et al*, 2015) was performed on polished particles, which were then further subjected to 3D classification without alignment to remove heterogeneity (Scheres, 2016). This homogenous set of particles was selected, 3D refined and post-processed. For calculating the final resolution, gold standard FSC = 0.143 criterion was used (Rosenthal & Henderson, 2003). Local resolution was estimated using Relion’s own implementation. The final map was auto-sharpened using Phenix.auto_sharpen (Terwilliger *et al*, 2018). Also, LocScale 2.0 was used to do local sharpening and to further improve map quality (Bharadwaj *et al*, 2026).

### Data processing of 50S-EngA•GTPγS complex

Initial 50S ribosome EngA•GTPγS data was processed in RELION 4.0.0. Reference-based picking was used to auto-pick 2,046,285 particles, and these particles were subjected to multiple rounds of 2D classification. For 3D classification, 1,258,447 particles were selected from the best 2D classes and classified into 10 classes. Classes 1, 3, 5, 7 and 8 show the association of EngA with 50S ribosomal subunit, while the rest of the classes contain unaligned particles. 642,131particles with 50S ribosome & EngA were selected and subjected to particle polishing to generate a consensus map with a resolution of 2.8 Å. To further remove the heterogeneity from the particles, 3D classification without alignment with T =20 was performed. The particles were then further classified into 10 classes, particles with 50S ribosome & EngA contributing to 312,230 particles were selected for another round of particle polishing and refinement. A 3D classification, into 5 classes, without alignment T = 200 was performed on polished particles (Supplementary Figure 2A). Further, to plot the angular distribution of the particles, angdist was used, and local resolution was estimated with Relion’s own implementation. The phenix.auto_sharpen was used to auto-sharpen the final cryo- EM maps (Supplementary Figure 2B).

Final **s**ingle-particle data processing for both datasets was performed using RELION 3.1.4. Data was collected in two sets in electron-counted movie mode. For Data 1 and Data 2, a total of 4812 and 4708 movie stacks were collected, respectively. A total of 969,121 and 762,509 particles were auto-picked and subjected to two rounds of 2D classification, yielding 470,158 and 438,618 particles for Data 1 and Data 2, respectively. These particles were subjected to 3D classification separately, and the classes containing EngA density with 50S ribosome, comprising 274,363 and 321,236 particles, were selected. These particles were then further refined and polished. The polished particles from both datasets were merged and refined to generate a consensus map with an overall resolution of 2.8 Å. To remove the heterogeneity, the particles were classified into 10 classes by using 3D classification without alignment and T = 20 (Supplementary Figure 3A). The best class with 272,928 particles was further selected and subjected to partial signal subtraction followed by focused 3D classification into 3 classes without alignment, to improve the cryo- EM density of EngA protein. The best class with 182,740 particles was selected; these particles were refined and post-processed to generate a final map with an overall resolution of 2.8 Å. The local resolution was calculated using Relion’s own implementation, phenix.auto_sharpen was used to auto-sharpen the final cryo- EM maps and angdist was used to plot angular distribution of the particles (Supplementary Figure 3B).

### Data processing of 50S ribosome EngA

After auto-picking, 2,084,989 particles were subjected to multiple rounds of 2D classification in RELION 3.1.4.. For 3D classification, 775,674 particles were selected from the best 2D classes, and these particles were divided into 10 classes. The classes in which EngA was associated with 50S ribosomal subunit were selected, which accounted for approximately 72% particles (537,759). These particles were then refined and polished, which yielded 3.0 Å resolution cryo-EM map. Further, 3D classification into 10 classes without alignment with T =40 was performed. Among these classes, one of the best class having 17.2% particles (92,830) with EngA associated with 50S ribosome was selected. Meanwhile, 3.3% of particles were mature 50S ribosome particles, 30% of particles were pre-50S ribosome, 31.8% of particles were 50S ribosome with H68-69 disordered, 14.8% of particles were 50S with immature CP and 2% of particles remained unaligned. Particles having EngA associated with 50S ribosome were then further subjected to another round of refinement and polishing to remove aberrations. These polished particles were then classified into three classes without alignment (T=20). Class 1 (24.4%) and class 3 (30.1%) particles showed distorted density for EngA, while class 2 (45.4%) particles have good density for EngA. Class 2 was further selected yielding 42,175 particles were refined and post-processed to generate map with an overall resolution of 3.4 Å. Relion’s own implementation was used to calculate the local resolution, auto-sharpening of the final cryo- EM map was done by phenix.auto_sharpen, and angdist was used to plot angular distribution of the particles (Supplementary Figure 4).

### Data processing of 50S-EngA•GDP

The Laplacian-of-Gaussian (LoG) filter was used to auto-pick initial 3,383,005 particles for 2D classification. After multiple rounds of 2D classification, 1,606,239 particles were further selected for 3D classification into 10 classes. Among the best 3D classes, 934,469 particles were selected, which were subjected to 3D refinement to generate a consensus map of 3.4 Å. These particles were then refined to remove aberrations and were further subjected to Bayesian polishing to remove beam-induced motion. These particles were then classified into 10 classes to remove the unaligned particles using 3D classification without alignment (T=20). Three classes showed the presence of EngA associated with 50S ribosome i.e, class 1 (74,619 particles), class 6 (129,113 particles) and class 8 (190,000 particles), all having different EngA conformation. Meanwhile, classes 2 and 10 were having empty 50S ribosome particles, classes 5 and 9 were pre-50S ribosome particles, and remaining classes contained unaligned particles. Further, high-resolution classes 6 and 8 were selected and separately refined to remove aberrations, followed by particle polishing (Supplementary Figure 5A). These polished particles from class 6 were then classified into three classes with blush regularization and without alignment (T=20). Classes 1 (60.4%) and 2 (1.2%) showed fragment density for EngA and unaligned particles, respectively. Moreover, class 3(38.4%) with good density for EngA was selected. These particles were refined with blush regularization and post-processed to generate a final map with an overall resolution of 3.3 Å (GDP S1.1). Similarly,, polished particles from class 8 were classified without alignment (T =20) and blush regularization into five classes. Among them, class 1 (60,523) and class 2 (129,761) particles had good density for EngA but different conformational states, while rest of classes had unaligned particles. These two classes were further refined with blush regularization separately. After post-processing of classes 1 and 2, final maps with overall resolution of 3.3 Å (GDP S2.2) and 3.2 Å (GDP S2.1) respectively, were generated (Supplementary Figure 5B). The resolution was estimated using the gold-standard FSC = 0.143 criterion, and local resolution was calculated using Relion’s own implementation. The phenix.auto_sharpen was used to auto-sharpen the final cryo- EM maps for classes 1 and 2, and angdist was used to plot the angular distribution of the particles (Supplementary Figure 5B).

### Data processing of 50S ribosome

The Laplacian-of-Gaussian (LoG) filter was used to select initial 617,170 particles for 2D classification. After multiple rounds of 2D classification, 496,303 particles were further selected for 3D classification into 10 classes. Among the best 3D classes, 334,076 particles were selected, which were subjected to 3D refinement to generate a consensus map of 3.9 Å. The refined particles were then classified into 10 classes without alignment. Classes 2, 9 and 10 with total of 169,483 particles were selected, while remaining classes showed pre-50S and low-resolution particles. These particles were again refined and classified into 5 classes without alignment to improve overall quality and resolution. Among these classes, class 1 (42.8%) shows the presence of mature 50S particles, class 3 (6.6%) shows pre-50 particles, classes 2 (6.6%) and class 5 (12.3%) show the presence of 70S particles, and class 4 (7.9%) particles contain unaligned particles (Supplementary Figure 56A). Classes 1 and 3 were further refined and post-processed to generate cryo- EM maps with an overall resolution of 3.0 Å and 3.2 Å, respectively. The local resolution was calculated using Relion’s own implementation, phenix.auto_sharpen was used to auto-sharpen the final cryo- EM maps, and angdist was used to plot angular distribution of the particles (Supplementary Figure 56B).

### Model building and structure analysis

The atomic coordinate of *M. smegmatis* 50S ribosome (PDB ID: 8WHY) (Kumar *et al*, 2024) was docked in the final cryo- EM map using ChimeraX (Goddard *et al*, 2018). The refinement was performed using phenix.real_space_refinement (Afonine *et al*, 2018). For EngA model building, the initial models were obtained from AlphaFold2 (Jumper *et al*, 2021) and docked in cryo- EM maps. The model building was carried out using COOT v.0.9.377 (Emsley *et al*, 2010). An iterative cycle of model building in COOT, followed by model refinement in PHENIX, was performed to model the rRNA helices whose conformations change upon EngA binding.

### Ribosome isolation from the *engA* knockdown and complemented strains

Freshly grown primary cultures of the *engA(-)*, *engA(-)*::*MSMEG_3738* and *engA(-)*::*Rv1713* strains were inoculated at an initial OD600 of ∼0.05 in 300 ml culture medium in the presence or absence of 100 ng/ml anhydrotetracycline (ATc). Cultures were pelleted after 24 hours of incubation at 37 °C with shaking at 200 rpm. The culture pellets were flash-frozen using liquid nitrogen. The ribosome was isolated following the same protocol, as described above (Scheres, 2012)(Goddard *et al*, 2018)(Emsley *et al*, 2010)

## Data availability

Cryo- EM maps and corresponding coordinates are being submitted to EMDB and PDB databases, respectively.

## Acknowledgements

We acknowledge the Electron Microscopy Facility at ATPC, RCB, Faridabad, for initial cryo-EM sample screening, and National Electron Cryo- Microscopy facility, BLiSc, C-CAMP-Bangalore, ANRF National Facility for Cryo- Electron Microscopy at IIT Kanpur, IIT Madras & SATHI, IIT Delhi, for cryo-EM data collection. We acknowledge the computing resource at the Indian Biological Data Centre (IBDC), Faridabad, used for data processing. We thank Prof. Umesh Varshney for valuable input during manuscript preparation and the P.S.K. lab members for proofreading the manuscript. We are grateful to Dr. William Jacobs at the Albert Einstein College of Medicine, USA, for providing us with the mycobacterial strain. A.A. & J.A. acknowledge the fellowship from the Department of Biotechnology (DBT/2021-22/RCB/1496) & DST-Inspire Govt. of India (DST/INSPIRE Fellowship/2022/IF220555), respectively. N. A. acknowledges funding support from the THSTI core research funding from the Department of Biotechnology, and P.S.K. acknowledges funding support from the ANRF CRG grant CRG/2022/002656 and the Regional Centre for Biotechnology, intramural grant.

## Author Contributions

P.S.K. & N.A. conceived the research project. A.A. purified the ribosomes and EngA protein, and reconstituted the ribosome-EngA complexes, collected cryo- EM data. J.A. generated the mutant clones, A.A. purified the mutants, and performed the sucrose pelleting assay. N.A. and J.A. generated the recombinant strains and performed the knockdown experiments. A.A. and P.S.K. performed the single particle reconstruction, model building, and structure analysis. P.S.K., A.A., N.A., and J.A. wrote the manuscript.

